# ClinePlotR: Visualizing genomic clines and detecting outliers in R

**DOI:** 10.1101/2020.09.05.284109

**Authors:** Bradley T. Martin, Tyler K. Chafin, Marlis R. Douglas, Michael E. Douglas

## Abstract

Patterns of multi-locus differentiation (i.e., genomic clines) often extend broadly across hybrid zones and their quantification can help diagnose how species boundaries are shaped by adaptive processes, both intrinsic and extrinsic. In this sense, the transitioning of loci across admixed individuals can be contrasted as a function of the genome-wide trend, in turn allowing an expansion of clinal theory across a much wider array of biodiversity. However, computational tools that serve to interpret and consequently visualize ‘genomic clines’ are limited. Here, we introduce the ClinePlotR R-package for visualizing genomic clines and detecting outlier loci using output generated by two popular software packages, bgc and Introgress. ClinePlotR bundles both input generation (i.e, filtering datasets and creating specialized file formats) and output processing (e.g., MCMC thinning and burn-in) with functions that directly facilitate interpretation and hypothesis testing. Tools are also provided for post-hoc analyses that interface with external packages such as ENMeval and RIdeogram. Our package increases the reproducibility and accessibility of genomic cline methods, thus allowing an expanded user base and promoting these methods as mechanisms to address diverse evolutionary questions in both model and non-model organisms.

## Introduction

Patterns of multi-locus differentiation, as distributed across admixture gradients, have long provided a window into divergence and speciation (e.g., Barton, 1983; Gompert, Mandeville, & Buerkle, 2017). Accordingly, they have been used to map loci associated with adaptation or reproductive isolation (Buerkle & Lexer, 2008; Martin et al., 2020), and as indicators of biotic responses to environmental change (Chafin, Douglas, Martin, & Douglas, 2019). Rather than relating these to patterns in the landscape, contemporary approaches have instead drawn conclusions based on genome-wide ancestries (Gompert & Buerkle, 2009; Fitzpatrick, 2013). The evolutionary processes that generate ‘genomic clines’ can be illuminated even when constituent taxa do not segregate geographically, but rather patchily (Bierne, Gagnaire, & David, 2013), or as a hybrid mosaic (Chafin et al., 2019).

Several programs are available specifically to investigate genomic clines. Of these, bgc (Gompert & Buerkle, 2011, 2012) is the most robust to false positives and uses a Bayesian approach that accounts for genotype uncertainty (Gompert, Lucas, et al., 2012) and autocorrelation caused by physical linkage (Gompert, Parchman, & Buerkle, 2012) in next-generation sequencing datasets. Although a powerful tool for analyzing hybridization with molecular data, it lacks user-friendly output. Researchers must either develop custom scripts or build cumbersome, one-off pipelines, neither of which is parsimonious. A more direct approach is clearly necessary.

Here, we present a comprehensive R-package, ClinePlotR, that promotes the genomic cline methodology. The package includes functions that facilitate bgc input file generation and output visualization and extend the plotting functionality from another genomic cline software package, Introgress (Gompert & Buerkle, 2010). Locus-wise clinal patterns are visualized by accessing a suite of R-methods that interpret them as a function of the genome-wide average, genomic position along chromosomes, and in relation to spatial and environmental parameters.

## Description

### Overall package workflow

The ClinePlotR R-package incorporates an introduction to available functions and can be installed via provided instructions directly from the GitHub repository (github.com/btmartin721/ClinePlotR). ClinePlotR includes three primary pipelines, a summary of which can be visualized in Figure 1.

**Figure 1:**
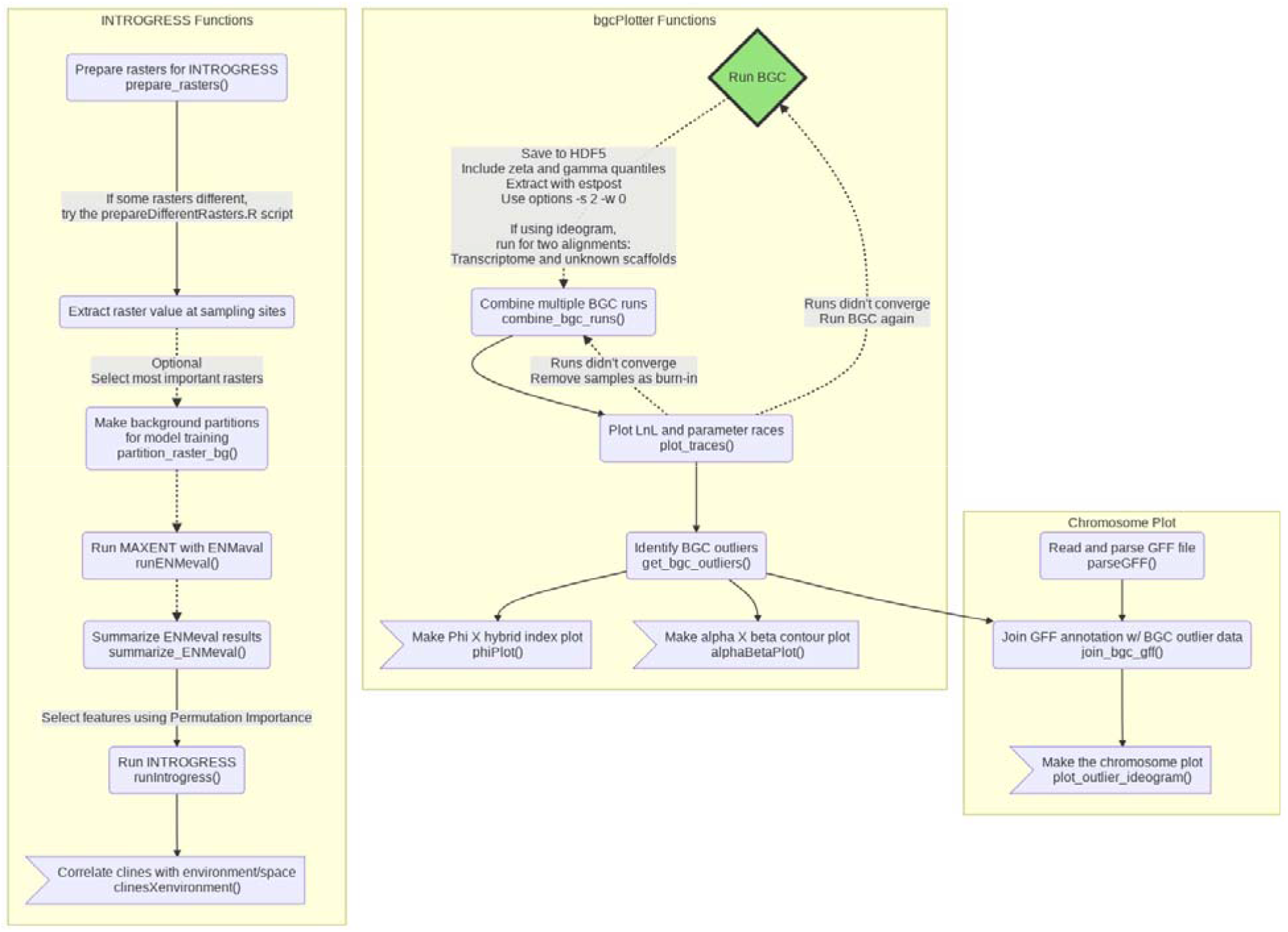
Simplified example workflow listing all available ClinePlotR functions. Yellow boxes group inter-dependent functions working towards producing one or two particular plots (terminal plotting steps depicted as flags). Connecting arrows indicate a pipeline where each step is dependent on the returned R objects. The green ‘Run BGC’ diamond identifies BGC as an external *a priori* step for the *bgcPlotter* and *chromosome plot* functions. The dotted lines indicate optional steps.

The workflow for our bgc pipeline includes functions to aggregate outputs from multiple independent runs, thin MCMC samples, and plot log-likelihood and bgc parameter traces. From these, ClinePlotR can both identify outlier loci using any of several user-defined options and plot locus-wise ancestry probabilities (ϕ) as a function of the hybrid index (Figure 2). Finally, users can examine the locus-wise relationship between cline center (α) and rate (β), with polygon hulls included to encapsulate 2D ‘outlier space’ for each parameter (Gauthier et al., 2020).

**Figure 2.**
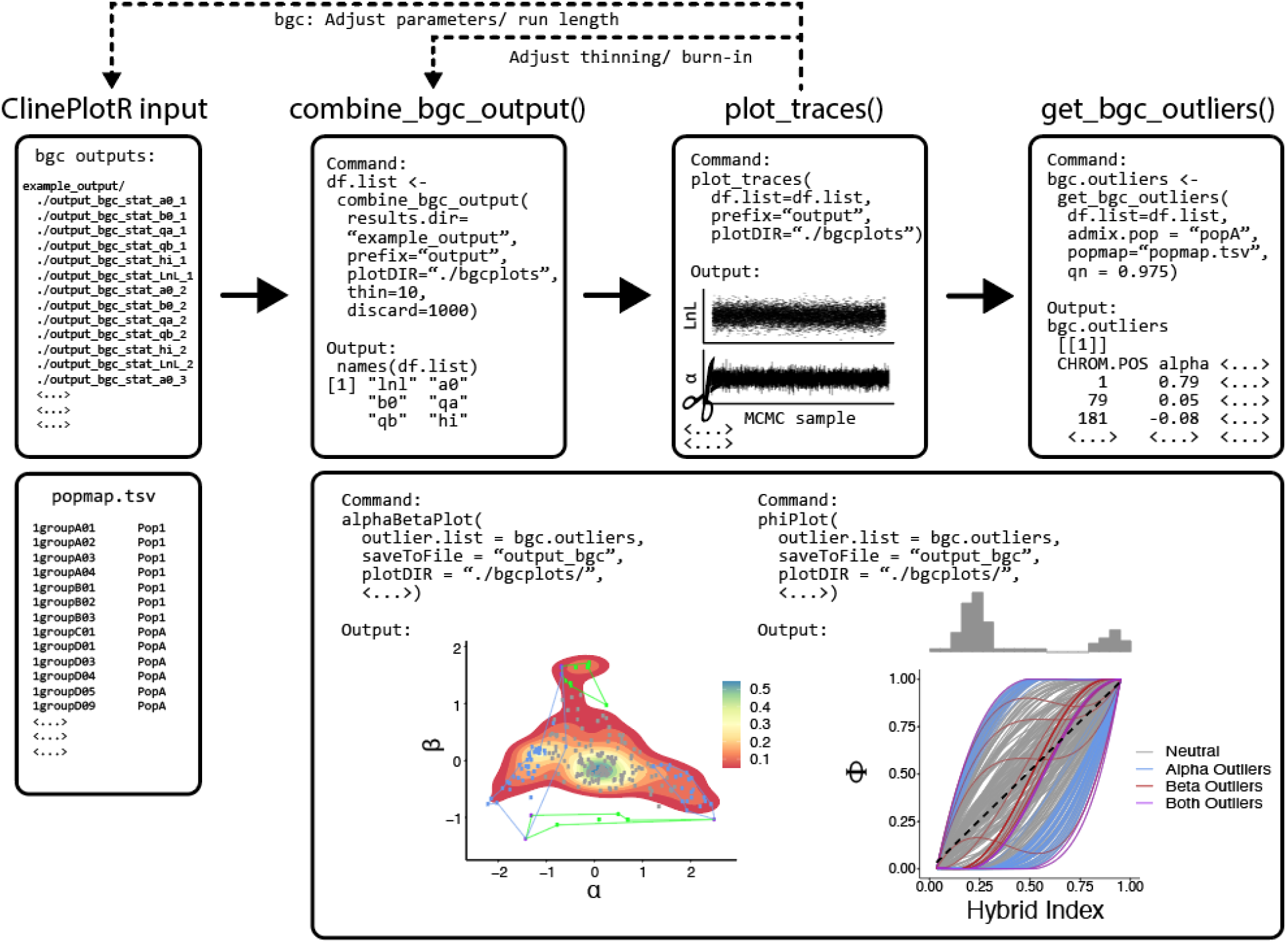
Example workflow for parsing Bayesian genomic cline (bgc) output, visualizing MCMC traces, detecting outliers, and plotting results. The ‘phiPlot’ (right-side, lower right box) shows hybrid indices (x-axis) and probability of parental population1 alleles (y-axis), plus a histogram of hybrid indices in the admixed population. The ‘alphaBetaPlot’ (left-side, lower right box) shows 2-D density of cline width/ rate representing the cline center (i.e., bias in SNP ancestry; α; x-axis) and steepness of clines (β; y-axis). Outliers are additionally encapsulated using polygon hulls.

ClinePlotR additionally includes accessory functions that allow an examination of variation in clinal parameters across the genome. Although mapping loci to reference assemblies is outside the scope of this package, an example of a workflow using minimap2 (Li, 2018) is in the documentation. If the user has access to physical SNP (single nucleotide polymorphism) coordinates and a closely-related chromosome-level assembly, ClinePlotR can integrate these data with the RIdeogram package (Hao et al., 2020) to yield karyotype-style ideograms annotated with heatmaps for both bgc cline parameters (Figure 3).

**Figure 3.**
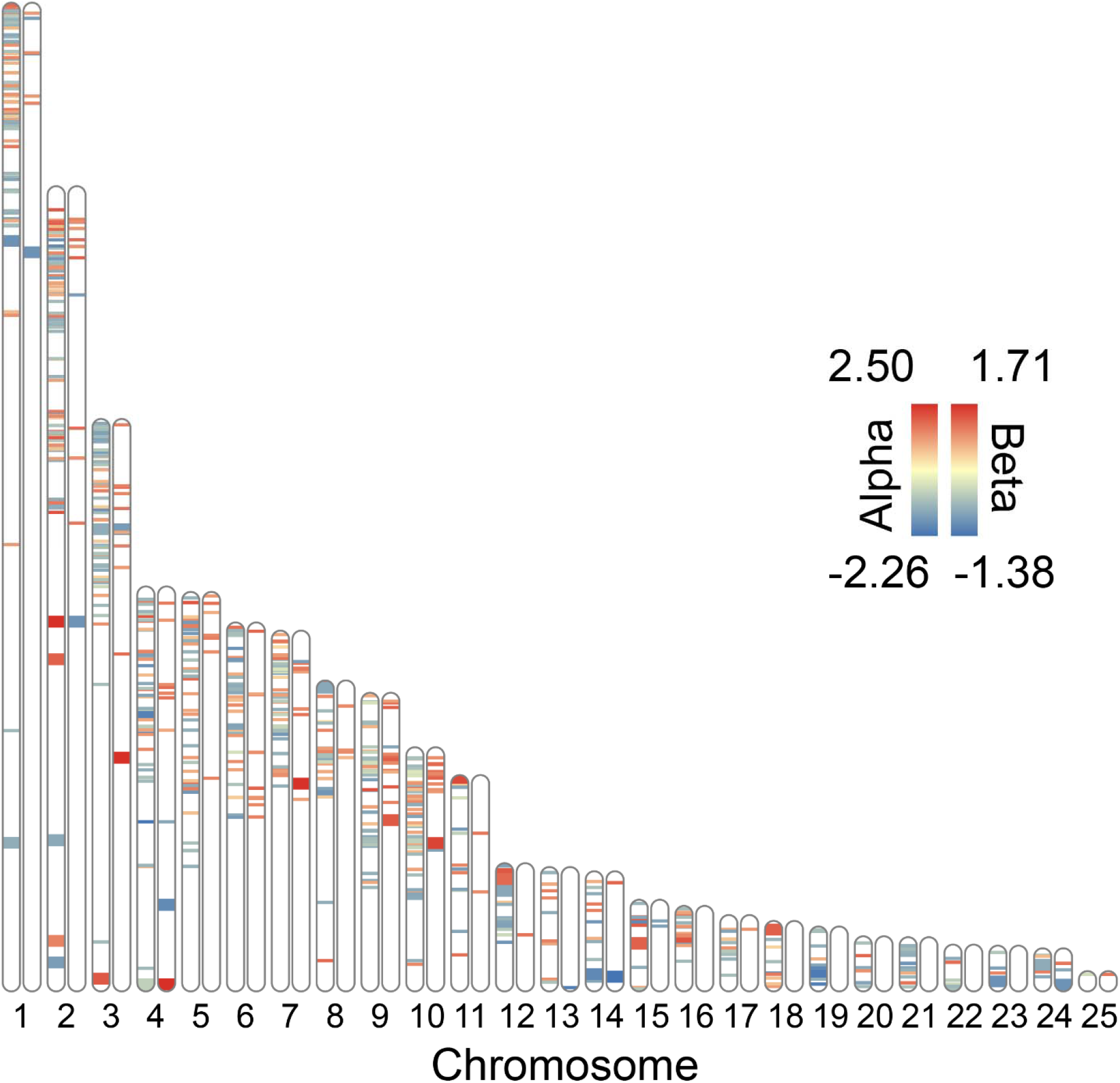
Example ideogram plot using Bayesian genomic cline (bgc) outliers for *Terrapene* ddRAD SNPs (y-axis), plotted onto *Trachemys scripta* chromosomes (x-axis). Chromosomes are duplicated, with alternative heatmaps for cline center (α; left) and rate (β; right). Larger heatmap bands correspond to SNPs located within known genes, whereas smaller bands were found in unknown scaffolds.

Functions are also provided to facilitate an Introgress workflow by generating input data frames as well as accessories that embellish the plotting functions already present in Introgress. These accessory functions will visualize spatial patterns (e.g., latitude/ longitude) and environmental variables that are inherent to genomic clines (Figure 4), to include helper functions that invoke ecological niche models (Maxent: Phillips, Anderson, & Schapire, 2006) as generated in the R-package ENMeval (Muscarella et al., 2014).

**Figure 4:**
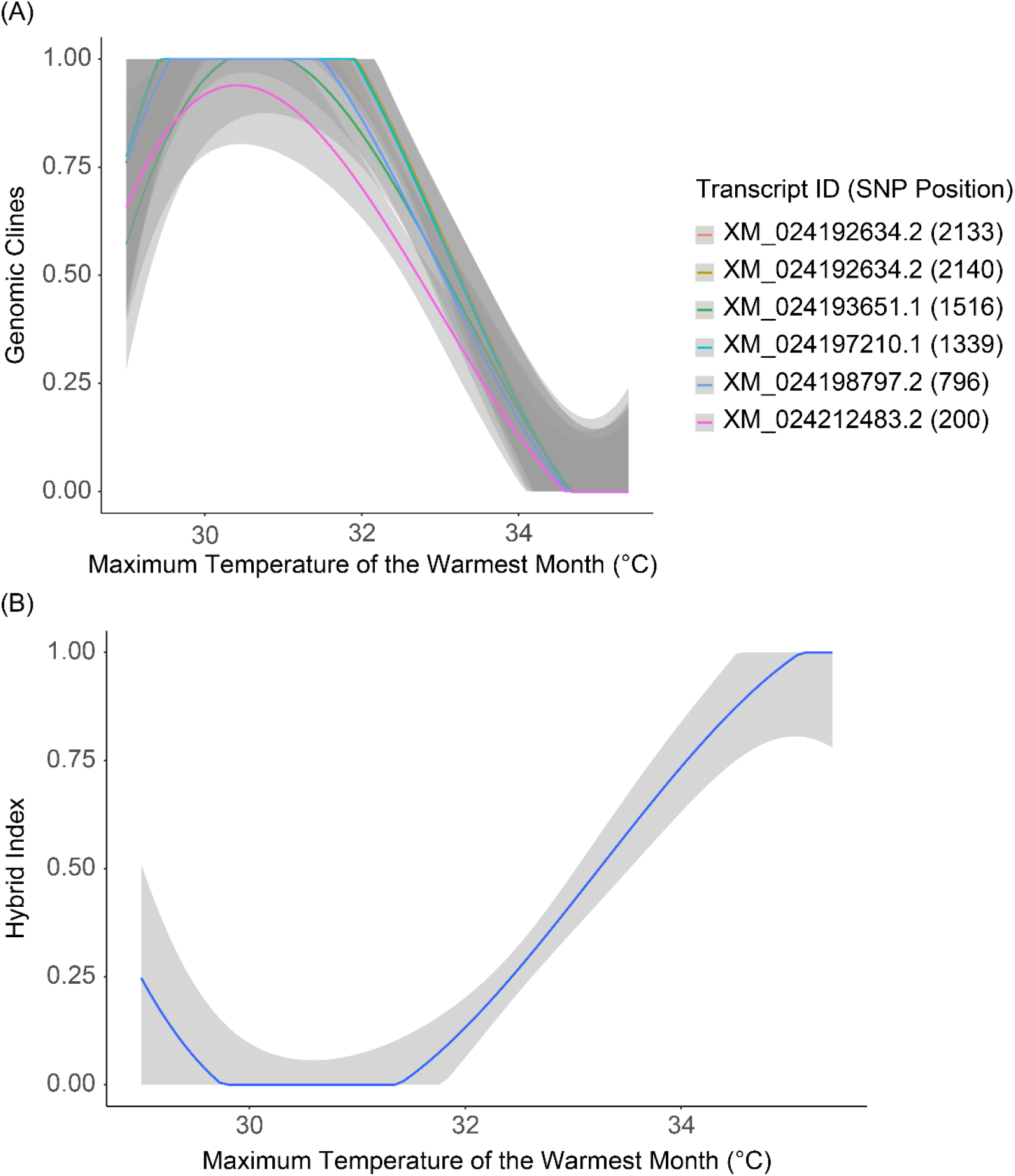
Example plots that can be made using the Introgress pipeline in ClinePlotR. The includes climatic variable on the X-axis corresponds to BioClim raster layer 5 (https://worldclim.org). The gray shading indicates confidence intervals for each regression line. (A) Genomic clines for six outlier SNPs mapped to the *Terrapene mexicana triunguis* transcriptome. Transcript IDs correspond to GenBank accession numbers and the position of each SNP (in base pairs) on the locus. (B) Hybrid index output from Introgress *versus* an environmental variable.

### Input and file format

The primary purpose of ClinePlotR is to simplify the use of rather cumbersome software designed to estimate genomic clines. To facilitate this task, accessory scripts that prepare files for input into bgc and Introgress are available in the GitHub repository, with a few variants. For example, *phylip2bgc.pl* script converts a PHYLIP-formatted alignment containing concatenated SNPs to the custom bgc input format. It can also subset populations and/ or individuals from a larger alignment. A similar script, *phylip2introgress.pl*, does likewise with Introgress input. Because bgc can additionally consider linkage among loci as well as genotype uncertainty, an input script (*vcf2bgc.py*) that employs the pyVCF Python library (https://pyvcf.readthedocs.io/) is also provided as a means to format an ipyrad (Eaton & Overcast, 2020) VCF file containing annotations for physical position and genotype read counts. Finally, an additional script, *nremover.pl*, is provided to comprehensively filter a PHYLIP-formatted SNP file. The program includes the capacity to filter by matrix occupancy per individual and per SNP column, and by minor allele frequency. It will also remove non-biallelic or monomorphic SNPs, and can randomly subsample large datasets.

### Outlier detection for Bayesian genomic clines

bgc output (extracted from HDF5 format using bgc’s *estpost* function) must be named as *prefix*_bgc_stat_*param*_*replicate*, where *prefix* is shared across all independent bgc replicates, *param* is an individual output parameter (e.g., LnL), and *replicate* is an integer. Outputs from any number of replicates can then be parsed, thinned, and combined via the *combine_bgc_output* function in ClinePlotR. The *combine_bgc_output* function provides arguments for the number of MCMC samples to be removed as burn-in, and for a sampling frequency with which to thin samples. Following bgc run aggregation, the MCMC samples can be visually inspected for mixing and convergence using a trace plotting function, *plot_traces*. Adjustments can then be made to thinning or burn-in parameters by re-running the *combine_bgc_output* function or, if necessary, by re-running bgc with altered parameters or increased MCMC length.

A primary goal of genomic cline analysis is to identify loci that possess either excess ancestry or exceptionally steep transitions relative to the genome-wide average. Here, we provide the function *get_bgc_outliers* that offers two outlier detection methods [described in Gompert & Buerkle (2011, 2012)]. Briefly, the first simply queries if the credibility intervals for the posterior probability distribution of cline parameters α or β (i.e., cline center and rate, respectively) exclude the neutral expectation (i.e., α or β = 0). If this interval excludes zero for either parameter, a locus can be flagged as either an α-outlier, β-outlier, or both.

The second method considers if per-locus parameter estimates are statistically unlikely, given the distribution of values across all loci. This is accomplished by classifying outliers as those for which posterior median α and β estimates are not encapsulated by the 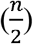 and 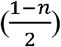 quantiles from a conditional α and β prior distribution (Gaussian with a mean of zero), where *n* represents a user-specified threshold (e.g., 95%, 97.5%). Users can choose whether to classify outliers using any combination of the above methods, but all require the zeta and gamma quantile estimates from the bgc output.

We additionally track whether parameter values are significantly positive or negative. This indicates either an increase (α > 0) or decrease (α < 0) in the probability of parental population ancestry among hybrids for a given locus, or deviation in the rate of transition in probabilities of locus-specific ancestries towards either very steep (β > 0) or wide (β < 0) shapes (Gompert & Buerkle, 2011).

### Visualization options

We attempted to tailor available visualizations in ClinePlotR towards common applications of Bayesian genomic clines found in the literature, and we will continue to add additional ones as need arises. Many applications seek to identify loci subject to various selective processes (Parchman et al., 2013) by comparing how ancestries transition among loci with respect to the genome wide average. To facilitate this, the *phiPlot* function computes ϕ_ijn_, the probability of parental population1 ancestry for each locus (*i*) and individual (*n*) within each admixed population (*j*) [Eqn. 3 and 4; Gompert & Buerkle (2011)]. It then produces a plot of ϕ (per locus) on the y-axis against posterior estimates of hybrid index on the x-axis (*sensu* Gompert et al., 2012), with a user-specified color scheme that designates statistical outliers (Figure 2).

Other applications have specifically examined relationships among cline rate and center parameters (Gauthier et al., 2020), and we also do so by implementing the *alphaBetaPlot* function. A 2-D density contour plot of α and β parameters is produced, with values for individual loci optionally mapped, and with the potential to calculate and plot polygon hulls that encapsulate positive and negative outliers with respect to each parameter (Figure 2).

### Extended functions and helper scripts

We also provide several additional functions (see Figure 1) that have considerable use cases, although some seemingly deviate from the ‘core’ bgc workflow. The first of several can be used to map parameter values of bgc clines onto a chromosomal ideogram via the function *plot_outlier_ideogram* (e.g., Figure 3), Here, bgc results are depicted for a case study examining hybridization between Woodland (*Terrapene carolina carolina*) and Three-toed box turtles (*Terrapene mexicana triunguis*) (Martin et al., 2020). However, some external user-steps are required to use the function.

Briefly, we mapped the *Terrapene* ddRAD sequencing alignment against the available *Terrapene mexicana triunguis* scaffold-level assembly (GenBank Accession: GCA_002925995.2). Scaffold coordinates were then converted to chromosome coordinates by mapping *Terrapene* scaffolds against the closely related chromosome-level *Trachemys scripta* assembly [(Simison, Parham, Papenfuss, Lam, & Henderson, 2020); GenBank accession: GCA_013100865.1]. This was accomplished by employing Minimap2 (Li, 2018) and PAFScaff (github.com/slimsuite/pafscaff). The output from *get_bgc_outliers* and PAFSCAFF, plus a GFF file read/ parsed via the provided functions *parseGFF* and *join_bgc_gff*, were used to plot a heatmap of bgc α- and β-values on an ideogram. Essentially, the ideogram plot (generated using the RIdeogram R-package) allows the chromosomal locations of each outlier to be visualized (Figure 3). It also provides a distinction between transcriptomic SNPs falling within known genes *versus* loci from surrounding scaffolds. For additional details, a more in-depth tutorial is provided at github.com/btmartin721/ClinePlotR.

Other extended functions include a wrapper to simplify running Introgress (*runIntrogress*), and a function that allows genomic clines (Figure 4A) and hybrid indices (Figure 4B) from Introgress to be correlated with spatial and environmental variables. To access this functionality, one can run *clinesXenvironment* using the object returned from *runIntrogress* and raster values extracted from each sample locality. Multiple rasters can be included (e.g., the 19 BioClim layers; https://worldclim.org/), and users can run the included ENMeval wrapper functions (*runENMeval* and *summarize_ENMeval*) to identify uninformative layers that may subsequently be excluded from *clinesXenvironment*. These latter functions access Maxent using the ENMeval pipeline (Muscarella et al., 2014), whereby the most informative raster layers are designated with the ‘permutation importance’ statistic.

## Conclusions

Genomic clines are useful for assessing patterns of introgression in hybrid zones. Unfortunately, parsing and plotting results from the available genomic cline software can be difficult. Given that genomic clines have a variety of applications, to include conservation genetics, evolutionary biology, and speciation research, it is clearly important that they be accessible for use by researchers. Here, we present an R-package that greatly simplifies the parsing of output from available genomic cline software, as well as the production of publication-quality figures. Our R-functions are intended to be user-friendly, and to this end employ a variety of parameters that can be altered by users to suit specific research needs. Furthermore, ClinePlotR allows outlier SNPs to be visualized across the genome, while also distinguishing known genes and surrounding loci. In addition, the environmental and spatial effects on genomic clines can be assayed. This extended functionality enhances the interpretation of genomic clines and provides greater insight into those underlying processes that potentially contribute to the observed patterns. Hopefully, future iterations of genomic cline software can act to extend chromosomal and environmental associations, particularly as whole genome sequencing becomes less expensive and more common.

## Acknowledgements

We thank S.M. Mussmann and M.R. Bangs for contribution to the *nremover.pl* script. We also thank staff of the Arkansas High Performance Compute Center (AHPCC) for access to computational resources that were used to test this package on the University of Arkansas Trestles cluster (funded through multiple National Science Foundation grants and the Arkansas Economic Development Commission).

## Authors’ contributions

BTM and TKC developed the R-package and wrote the manuscript and all code. MRD and MED were the study supervisors. All co-authors contributed to editing the manuscript.

## Data accessibility

ClinePlotR is available as a GitHub repository: https://github.com/btmartin721/ClinePlotR. We also plan to submit to CRAN prior to publication. The data used herein will be available as an example dataset in a Dryad Digital Repository [DOI]. During review, the data will also be temporarily accessible from Box Drive (https://uark.box.com/s/ei21v3unvxfxczbfnfflwhrk536bsd5h)

